# Polyglutamine-related aggregates serve as a potent antigen source for cross presentation by dendritic cells

**DOI:** 10.1101/806414

**Authors:** Shira Tabachnick-Cherny, Dikla Berko, Sivan Pinto, Caterina Curato, Yochai Wolf, Ziv Porat, Rotem Karmona, Boaz Tirosh, Steffen Jung, Ami Navon

## Abstract

Protective MHC-I dependent immune responses require an overlap between repertoires of proteins directly presented on target cells and cross-presented by professional antigen presenting cells (APC), specifically dendritic cells (DCs). How stable proteins that rely on DRiPs for direct presentation are captured for cell-to-cell transfer remains enigmatic. Here we address this issue using a combination of *in vitro* and *in vivo* approaches involving stable and unstable versions of ovalbumin model antigens displaying DRiP-dependent and -independent antigen presentation, respectively. Apoptosis, but not necrosis of donor cells was found associated with robust p62-dependent global protein aggregate formation and captured stable proteins permissive for DC cross-presentation. Potency of aggregates to serve as antigen source was directly demonstrated using polyglutamine-equipped model substrates. Collectively, our data implicate global protein aggregation in apoptotic cells as a mechanism that ensures the overlap between MHC-I epitopes presented directly or cross-presented by APC and demonstrate the unusual ability of DC to process stable protein aggregates.

**Summary:** Protective T cell immunity relies on the overlap of the antigen repertoire expressed by cells and the repertoire presented by dendritic cells that are required to trigger naïve T cells. We suggest a mechanism that contributes to ensure this antigenic overlap. Our findings demonstrate that upon apoptosis stable proteins are aggregated in p62-dependent pathway and that dendritic cells are capable to efficiently process these aggregates to retrieve antigens for T cell stimulation.

## Introduction

The immune system evolved to sense and eliminate infected and transformed cells, taking advantage of peptide presentation on MHC class I (MHC-I) to trigger antigen-specific cytotoxic T lymphocytes (CTL) (1). MHC-I is expressed by all nucleated cells and generally presents peptides originating from intracellular proteins (2, 3). The MHC-I peptide repertoire is hence dynamic, reflecting the constant alteration of the intracellular proteome. This ‘direct presentation’ pathway samples the general peptide pool generated by proteasomal degradation, a peptide subset that is actively transported into the endoplasmic reticulum (ER) by the ATP-dependent transporter associated with antigen processing (TAP) (4). In the ER, peptides are loaded onto newly synthesized MHC-I molecules in a facilitated manner, and peptide-MHC-I complexes are transported for presentation to the cell surface.

MHC-I peptides can originate from various sources: short-lived proteins that are directed to the proteasome by regulated ubiquitination (5) and retirees of long-lived proteins (6). A third major MHC-I peptide source is believed to consist of defective ribosomal proteins (DRiPs) (7, 8). DRiPs arise through faulty initiation from downstream codons, premature termination, short protein segments produced by pioneer translation rounds during nonsense mediated decay (NMD), mRNA quality control (QC) and out-of-frame translated protein segments (9–12).

At the cell surface, MHC-I peptide complexes are constantly surveyed by CD8^+^ T lymphocytes carrying epitope-specific T cell receptors (TCRs) (13). Once complexes are recognized, CD8^+^ T cells are activated and acquire cytotoxicity that allows them to eliminate peptide-presenting specific target cells (14). The initiation of CTL responses against infected cells requires ‘priming’ by professional antigen presenting cells (APC) through an antigen cross-presentation pathway (15, 16)(Carbone et al., 1998; Jung et al., 2002), which is restricted to a BatF3-dependent CD8α^+^ dendritic cell (DC) subset (DC1) (17, 18). Cross-presentation requires endocytosis of cellular material, protein transport from the lysosome to the cytoplasm and proteasomal processing (19, 20). In contrast to direct presentation, cross-presentation hence relies in addition to proteasome activity (21) on prior proteolysis in the lysosome (22, 23). Polypeptides are delivered to the proteasome from the endocytic compartments, perhaps by forming ER/endosome mixed vesicles (24). Yet, the exact cascade of events that interconnects cytosolic and endocytic proteolysis for cross-presentation remains incompletely understood (25–29).

Efficient protective MHC-I dependent immune responses require the overlap between the antigenic MHC-I repertoires that are presented directly on infected cells and cross-presented by the APC (30). Biochemical mechanisms that ensure this repertoire overlap are poorly understood. Specifically, it is unclear how extremely stable proteins, such as virus-derived structural peptides (7), that rely for direct presentation on DRiPs, are captured for cross-presentation, as DRiPs are unavailable for cell-to-cell transfer.

Here we show that both stable and unstable variants of the model antigen ovalbumin (RFP-Ova & Ova-RFP) contribute antigenic epitopes for direct presentation in DRiP-dependent and DRiP-independent manner, respectively. Despite their distinct stabilities, both fusion proteins also performed equally well in cross-presentation of apoptotic cells, including pattern and magnitude of elicited *in vivo* responses, as well as their ability to induce functional T cell cytotoxicity. Non-selective aggregation that was induced in apoptotic, but not necrotic cells, prior to their engulfment by DCs, was found to constitute a major antigenic source for MHC-I cross presentation. Congruent with this notion, polyQ (94Q)-Ova derivatives, engineered to aggregate independently of apoptosis, served as potent long-lasting source for cross-presentation. Collectively, our data underscore protein aggregation as a major factor that contributes to the repertoire overlap between direct and cross-presented MHC-I antigens of stable cellular proteins.

## Results

### DRiP-dependent and -independent MHC class I antigen presentation of OVA model antigens

To study mechanisms regulating the overlap between direct and indirect MHC-I presentation we engineered two proteins of distinct stability that share identical primary sequence; both substrates included the intact ovalbumin (Ova) sequence fused to a stable globular monomeric red fluorescent protein (RFP), either at its N’ or C’ terminus (RFP-Ova and Ova-RFP, respectively). Studying proteasomal processing of these fusion molecules in EL4 cells, we previously demonstrated that ovalbumin was degraded only when its N-terminus was accessible (Ova-RFP) (31). When RFP was attached to the ovalbumin N’ terminus (RFP-Ova), the cellular abundance of the protein increased relative to Ova-RFP, despite similar mRNA levels (Figure 1A and 1B). This difference was attributed to proteasomal degradation, as Ova-RFP was stabilized in the presence of the proteasome inhibitor (Velcade) (Figure 1A).

**Figure 1:**
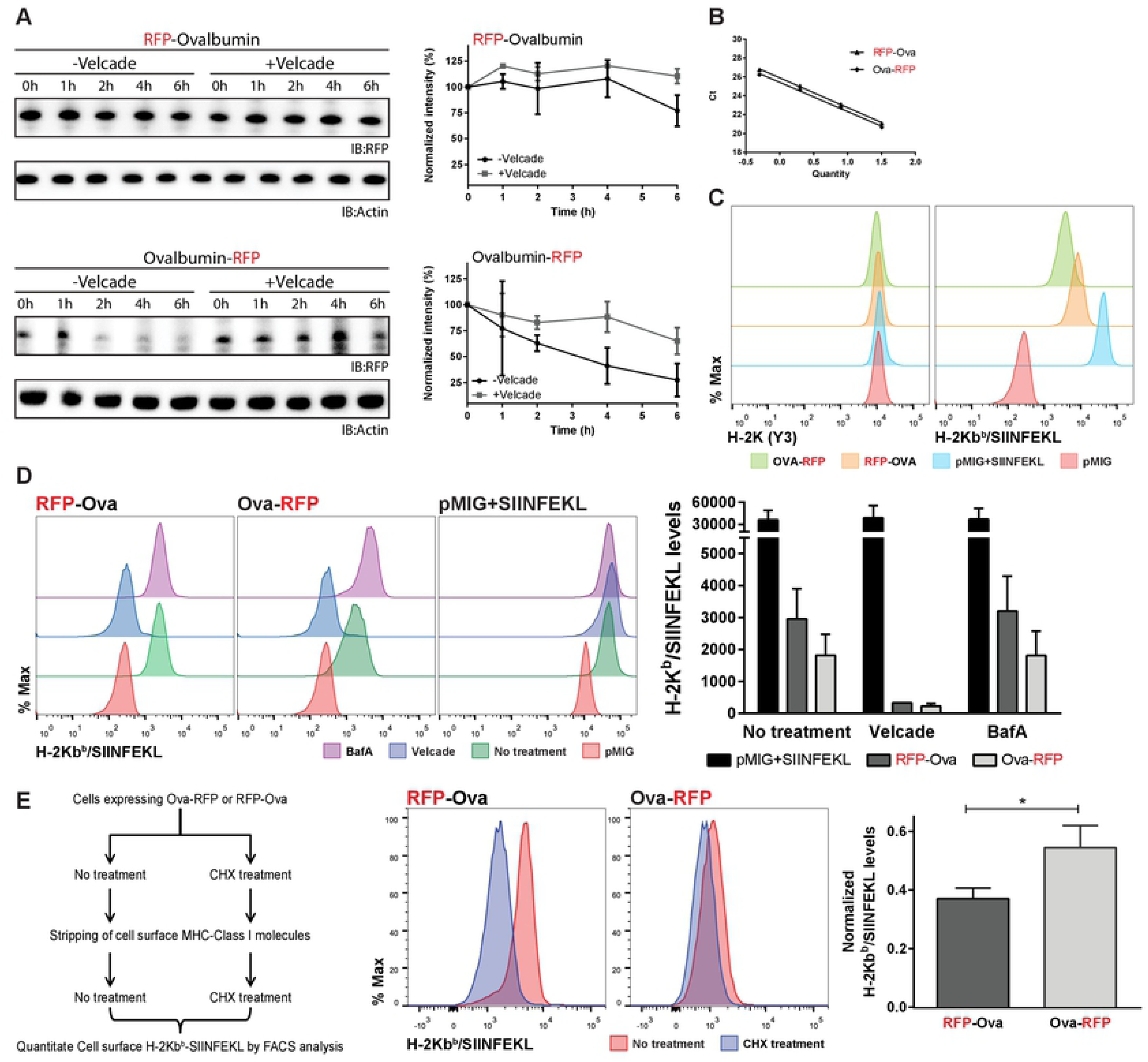
SIINFEKL is efficiently presented on MHC class I upon expression of short-lived Ova-RFP and long-lived RFP-Ova derivatives. **(A)** Western blot analysis of transfected EL4 cells constitutively expressing RFP-Ova or Ova-RFP. Cells were pre-incubated at 37°C in presence and absence of 2µM of Velcade (Bortezomib, proteasome inhibitor) for an hour. Next, CHX was added (50 μg/ml) for the indicated times. Cell lysates were subjected to Western blot analysis by anti-RFP and anti-actin antibodies. Quantification of the residual amount of protein detected at each time point during the chycloheximide chase is presented (SD were calculated from three repetitions under similar conditions). **(B)** qPCR calibration curve from cells expressing either RFP-Ova or Ova-RFP demonstrating similar mRNA levels. **(C)** Flow cytometry analysis of transfected EL4 cells expressing either pMIG vector (plasmid without OVA variant) (negative, red), pMIG cells loaded with 10ug/ml SIINFEKL (light Blue), pMIG RFP-Ova cells (orange) or pMIG Ova-RFP cells (green). Cells were labeled with either H-2K antibody (Y3 hybridoma - left) or H-2Kb^b^/SIINFEKL antibody (25D1.16 hybridoma - right). **(D)** Flow cytometry analysis of transfected EL4 cells expressing either pMIG vector (negative, red), Ova-RFP/RFP-Ova or pMIG+SIINFEKL (positive control, green) comparing the effect of either Velcade, proteasome inhibitor (blue) or Bafilomycin A, lysosome inhibitor (purple) on H-2Kb^b^/SIINFEKL levels. Cells were incubated for two hours at 37°C with the relevant inhibitors, then MHC class I-bound peptides were stripped of the cell surface by acid wash and the cells were incubated with or without the indicated inhibitor for additional 6 hours. Cells were stained with 25D1.16 antibody. Statistics from 3 independent experiments are demonstrated in the graphs. **(E)** Left lane: Schematic flow chart of the experiment. Mid histograms: Flow cytometry analysis of EL4 cells, expressing either Ova-RFP or RFP-Ova, incubated with or without 50µg/ml CHX at 37°C, as illustrated at the schematic flow chart. Cells were stained with 25D1.16 antibody that recognizes H-2Kb^b^/SIINFEKL complexes. Statistics from 5 independent experiments are presented on the left graph, with significance of p<0.05 (*). Right graph demonstrates the normalized values of H-2Kb^b^/SIINFEKL with the general levels of H-2K (Y3 antibody), statistics derived from 3 independent experiments.

Cells expressing either RFP-Ova or Ova-RFP were examined for their steady state H-2K^b^ MHC-class I levels and their specific MHC/peptide complex display (Figure 1C). The immuno-dominant Ova MHC-I epitope in C57BL/6 mice (SIINFEKL) was efficiently presented on EL4 cells expressing either RFP-Ova or Ova-RFP, as indicated by antibody surface staining for H-2K^b^/SIINFEKL complexes (32) (Figure 1C). Sensitivity of presentation to Velcade, but resistance to the lysosomal inhibitor Bafilomycin A (BafA) confirmed that SIINFEKL presentation was proteasome-dependent for both Ova-RFP and RFP-Ova expressing cells (Figure 1D). Notably, H-2K^b^ surface levels were moderately affected by Velcade treatment, but not BafA (Supplementary Figure S1), and the effect did not reach significance. All flow cytometry analyses were controlled for cell viability using Annexin V and DAPI staining (Supplementary Figure S2). Taken together, these findings suggest that Ova epitopes detected on cells expressing RFP-Ova derive mainly from ribosome-associated quality control (QC), rather than the pool of stable RFP-Ova molecules. To further substantiate this notion, RFP-Ova and Ova-RFP expressing EL4 cells were incubated with or without the translation inhibitor cycloheximide (CHX) and H-2K^b^/SIINFEKL levels were examined. CHX treatment resulted in a larger decrease of surface complexes in cells expressing stable RFP-Ova compare to cells expressing the unstable Ova-RFP (Figure 1E). These data are consistent with a higher contribution of ribosome-associated degradation, i.e. DRiPs, as a source of MHC-I epitopes of stable proteins than less stable proteins.

We conclude that DRiP-dependent (RFP-Ova) and -independent (Ova-RFP) model antigens constitute a valuable system to elucidate mechanisms ensuring effective cross-presentation of epitopes that depend for direct MHCI presentation on QC processes.

### Cross-presentation efficacy of stable and unstable proteins match the efficiencies observed in direct presentation

Following CD8^+^ T cell differentiation into effectors, CTLs recognize and eliminate infected or transformed cells that display cognate antigen epitopes. Induction of effector functions requires specialized APCs, such as classical XCR1^+^ DCs (cDC1), that ingest apoptotic infected or transformed cells and shuttle proteins of the latter into their own MHC-I pathway (17). To test the efficiency by which RFP-Ova and Ova-RFP are captured for cross-presentation, we applied *in vitro* and *in vivo* T cell activation assays. The magnitude of *in vitro* T cell activation directly correlated with the level of H-2K^b^/SIINFEKL complexes presented on cells expressing either RFP-Ova or Ova-RFP (Supplementary Figure S3). The B3Z T cell hybridoma line expressing β-galactosidase under control of the IL-2 promoter showed a small difference in SIINFEKL presentation and the related T cell response in favor of RFP-Ova, which did not reach statistical significance (Supplementary Figure S3).

To measure *in vivo* T cell responses to cells expressing the OVA variants, we engrafted C57BL/6 wild type (WT) animals with H-2K^b^/SIINFEKL-reactive T cells isolated from TCR transgenic OT-I mice (33). Grafted cells expressed an allotypic marker (CD45.1) to distinguish them in the recipients and were labeled with the intracellular dye CarboxyFluorescein Succinimidyl Ester (CFSE) to monitor their proliferation (34). Engrafted mice were immunized with apoptotic cells expressing the Ova derivatives (Figure 2A). Cells expressing either of the different OVA constructs or no Ovalbumin, displayed comparable apoptotic cell percentages (Supplementary Figure S4). Flow cytometric analysis of lymph node and spleen cell preparations of the mice for OT-I T cells revealed robust T cell proliferation in response to both RFP-Ova and Ova-RFP immunizations, as indicated by CSFE dilution (Figure 2C). However, as opposed to the above *in vitro* experiment (Supplementary Figure S3), we observed a significant statistical difference between animals immunize with the two constructs in favor of RFP-Ova (Figure 2C).

**Figure 2:**
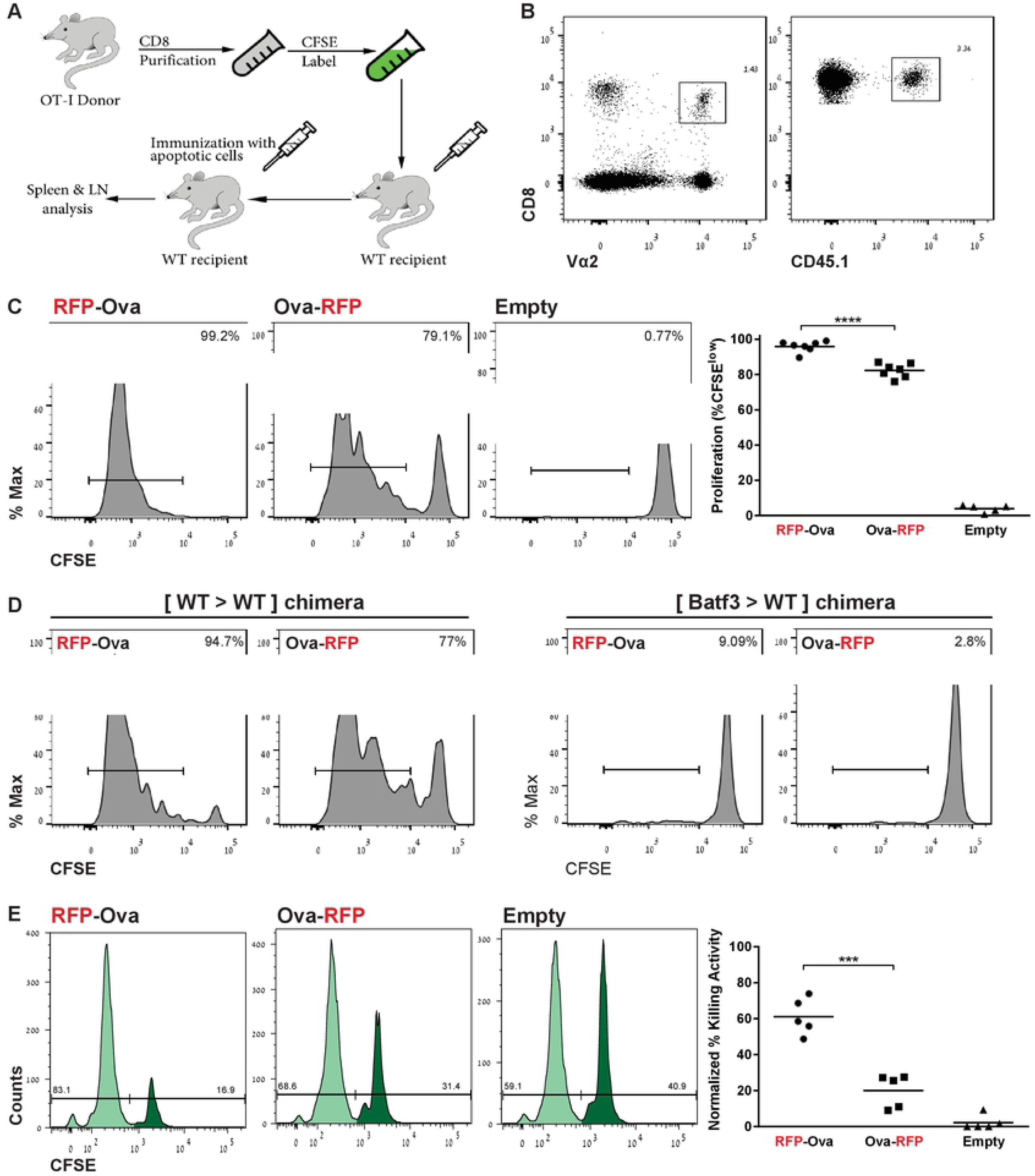
OT-I T cell activation and cytotoxicity correlate with MHC-I presentation levels of the different Ova derivatives. **(A)** Schematic of experimental protocol for (B & C). **(B)** Flow cytometric analysis of lymph node and spleen cells indicating gating for engrafted T cell population in CD45.2^+^ wt recipient. T cell graft is defined by expression of CD8, Vα2 and CD45.1. **(C)** Flow cytometric analysis of lymph node and spleen cells for proliferation (CFSE dilution) of CD8^+^Vα2^+^CD45.1^+^ T cell graft. Statistical significance is p < 0.0001 (****). Results are representative of three independent experiments, total n=7 (RFP-Ova), n=7 (Ova-RFP) and n=5 (Empty). **(D)** Flow cytometric analysis of splenocytes of immunized [WT > WT] and [*Batf3^-/-^* > WT] BM chimeras for CFSE dilution of CD8^+^Vα2^+^CD45.1^+^ T cell graft. n=10 [WT > WT] and n=12 [*Batf3^-/-^* > WT] chimeras. **(E)** In vivo killing assay (experimental scheme is provided in supplementary Figure S6). Flow cytometric analysis of spleens of immunized mice injected with SIINFEKL-loaded and control target splenocytes (CD45.1^+^), labelled with high CFSE (dark green) and low CFSE (light green), respectively. Histograms represent cell amount in each CFSE population; the ratio between the populations was calculated as described in the Experimental Procedures. On the right, a graphic summary of the calculated specific killing efficiency (Statistical significance of p< 0.001 is indicated by ***). Results are representative of two independent experiments, total n=5 for each RFP-Ova, Ova-RFP and Empty cells.

Development of the cDC1 DC subset dedicated to cross-presentation depends on the transcription factor BATF3 (18). To investigate if cDC1 are also required for the T cell priming observed in our system, we generated bone marrow (BM) chimeras by transplanting irradiated WT animals with Batf3-deficient (*Batf3^-/-^*) BM (18). Absence of cDC1 from spleens of [*Batf3^-/-^* > WT] chimeras was confirmed by flow cytometric analysis (Supplementary Figure S5). As opposed to [WT > WT] controls, proliferation of adoptively transferred OT-I cells was completely abrogated in immunized Batf3 KO chimeras (Figure 2D), establishing that processing of the Ova derivatives and cross-presentation of their SIINFEKL epitopes is executed predominantly by cDC1s.

For immune surveillance, MHC-I directed immune responses are meant to remove cells decorated with pathogen- or transformation-associated non-self epitopes. To test whether the immunizations with apoptotic cells expressing the Ova derivatives result in cytotoxicity, we used an *in vivo* killing assay (35). Immunization with apoptotic cells expressing the stable RFP-Ova that displayed superior presentation level of SIINFEKL and T cell stimulation also generated the most efficient *in vivo* killing activity (Figure 2E, Supplementary Figure S6). In contrast, immunization with Ova-RFP expressing apoptotic cells induced significantly less cytotoxicity (Figure 2E). Taken together, for the antigenic system we tested including stable and unstable model proteins, the efficiency of direct presentation matched the efficiency of cross-presentation, which is an important factor in inducing the ultimate protective CTL response.

### Global protein aggregation during apoptosis captures the stable cellular steady state proteome

As shown above, direct MHC-I presentation of RFP-Ova relies on DRiPs or other QC pathways. Arguably, these mechanisms provide insufficient protein material for transfer and cross-presentation by APCs. To directly address this issue, we monitored the abundance and the distribution of both Ova derivatives following apoptosis induction. Protein distributions in supernatant and pellet fraction were assessed as described previously (36). Briefly, cells were lysed in the presence of 1% NP-40 and the lysate was overlaid on a 20% glycerol cushion. Samples were centrifuged at 100,000g for 45 min at 4°C. As expected, in untreated cells, both the stable long-lived RFP-Ova derivative and the short-lived Ova-RFP derivative were detected almost exclusively in the supernatant (Figure 3A). In contrast, upon apoptosis induction, most of the proteins were detected in the insoluble pellet fraction, suggesting that apoptosis is accompanied by aggregate formation (Figure 3A). Notably, aggregation was not ensued upon cell synchronization, but only following treatment with etoposide (Supplementary Figure S7). Lamin B1 was used to assess the supernatant / pellet separation (Figure 3A), as lamin B1, a major component of nuclei, is known to appear in the insoluble fractions (37, 38). Notably, lamin B1 is processed upon apoptosis by caspase 7 and thus appeared truncated upon apoptosis providing a convenient additional internal control (39–42).

**Figure 3:**
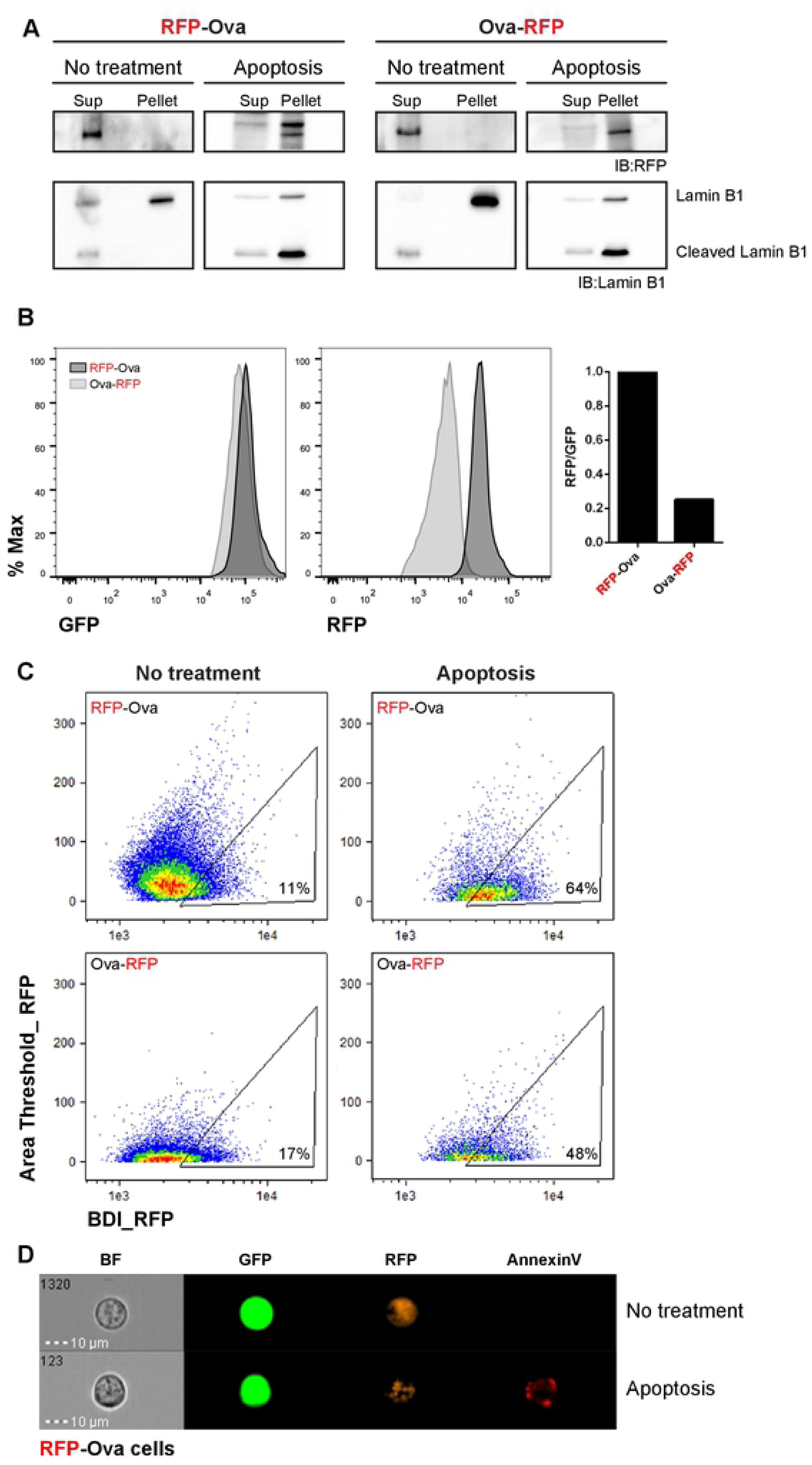
Aggregation during apoptosis captures the steady state proteome and directs MHC-I cross presentation. **(A)** Western blot analysis of transfected cells expressing Ova-RFP or RFP-Ova derivatives. Cells were treated for 12h with 500µM hydroxyurea following 12h of incubation with 100 µM etoposide. Equal numbers of cells were lysed and supernatant and pellet fractions were separated on a 20% glycerol cushion and analyzed with anti-RFP and anti-laminB1 antibodies. Results are representative of 3 independent repeats. **(B)** Flow cytometric analysis of 39.5 cells for GFP and RFP intensities representing transcription levels and abundance of Ova-RFP and RFP-Ova, respectively. Representative result of 2 repeats. **(C)** Image stream analysis of 39.5 cells before and after apoptosis induction by etoposide treatment. Apoptotic cells presented were gated for annexin V positivity. High BDI and low area threshold indicates aggregated RFP. Representative cells are displayed in **(E)**. Results are representative of two independent experiments. **(D)** Representative image demonstrating RFP-Ova distribution in 39.5 cells prior and following apoptosis induction, which was confirmed using annexin 5 labeling.

To quantify the extent of aggregation we performed a multispectral imaging flow-cytometry (ImageStream) analysis. While results of an IRES-GFP expression reporter assay indicated comparable transcript levels of the constructs, as indicated by identical GFP intensity (Figure 3B, left), amounts of the stable RFP-Ova protein were significantly higher than those of Ova-RFP, in accord with the short half-life of the latter (Figure 3B, right). Next, we determined the aggregation patterns of the two Ova derivatives during early apoptosis (Figure 3C-D) by assessing Bright Detail Intensity (BDI) and area threshold (43). BDI levels reflect the intensity of localized bright spots within the selected cell, which can serve as indicators of aggregates. In contrast, the area threshold refers to pixels, which illuminate in an intensity, which is within the highest 50% of the relative intensity scale of the measured sample. Accordingly, high values of area threshold indicate an even distribution of fluorescent proteins in the cell, while low values indicate uneven distributions.

Apoptotic cells were labeled with Annexin V and the positive (apoptotic) population was gated. As shown in Figure 3C, before treatment (NT) RFP-Ova expressing cells displayed low BDI and acquired higher BDI values upon apoptosis, indicating aggregates accumulation. A similar effect, yet less pronounced, was detected in Ova-RFP expressing cells undergoing apoptosis (Figure 3C). Representatives of RFP-Ova expressing cells prior and following apoptosis are shown in Figure 3D and Supplementary Figure S8, demonstrating the difference of RFP distribution.

We next tested if aggregate formation is uniquely associated with apoptosis, or also triggered by necrosis. Apoptotic and necrotic RFP-Ova expressing cells (induced by etoposide and heat shock respectively) were identified by AnnexinV (AV) and DAPI staining. Apoptotic cells demonstrated 25-30% of AV^+^DAPI^-^ and AV^+^DAPI^+^ populations. In contrast, necrotic cells did not comprise AV^+^DAPI^-^ events (0.62%) and the majority of the cells were AV^+^DAPI^+^ (86%) (Figure 4A), corroborating an earlier report (44). ImageStream analysis of the two populations for RFP levels and extent of aggregate formation revealed that global RFP fluorescent signals were similar in both populations (Figure 4B). In contrast, the fluorescent signal appeared diffused in necrotic cells, while accumulated in aggregates in apoptotic cells (Figure 4C and Supplementary Figure S8). In line with this observation, high BDI values were only detected in the double positive AV^+^DAPI^+^ apoptotic cells population (Figure 4D). DNA condensation in the cells was monitored by ‘area threshold and contrast parameters’ (Figure 4E). Apoptotic nuclei yielded small, fragmented, highly textured nuclear images with Draq5 dye localized to small punctate regions of the fragmented nucleus (45). In contrast, such high contrast nuclear staining was absent from the necrotic cell preparation. The distinction between apoptosis and necrosis was also verified biochemically. As demonstrated above, lamin B1 is processed upon apoptosis (Figure 3A) (42). In contrast, processing of lamin B1 was not detected in the necrotic cells preparation (Figure 4F). Taken together our findings suggest that in our system, apoptotic cell death, but not necrosis, is associated with global cellular protein aggregation.

**Figure 4:**
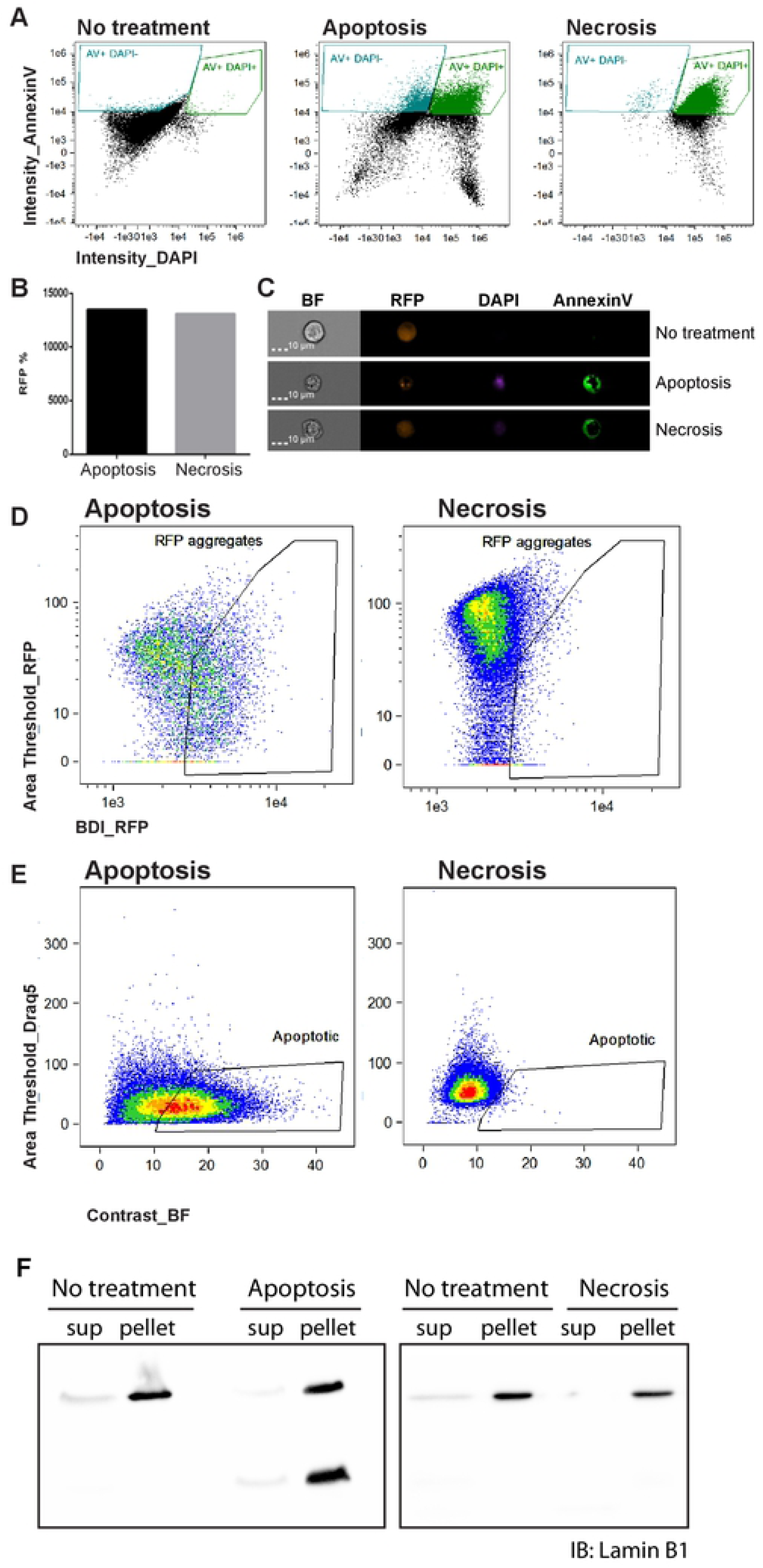
Aggregate formation occurs in cells undergoing apoptosis induction, but not in cells triggered for necrosis. **(A)** Image stream analysis of 39.5 cells expressing RFP-Ova construct, following apoptosis or necrosis induction by etoposide or heat shock, respectively. Cells were labeled with AV and DAPI and population distributions were analyzed. **(B)** Flow cytometric analysis of RFP intensity in necrotic and apoptotic AV^+^DAPI^+^ population. (**C**) Representative image demonstrating RFP-Ova distribution in 39.5 cells triggered for apoptosis or necrosis. (**D, E**) Image stream analysis of 39.5 cells indicating High BDI, low area threshold as well as high bright field contrast (E) revealing aggregated RFP, within the AV^+^DAPI^+^ gated populations. **(F)** Western blot analysis of transfected cells expressing Ova-RFP upon induction of apoptosis or necrosis. Cells were lysed, supernatant and pellet fractions were separated on SDS-PAGE and western blotted with anti-lamin B1 antibodies. Results are representative of 2 independent repeats. Similar results were obtained when RFP-Ova expressing cells were used (data not shown).

### p62 is involved in apoptosis related aggregates formation

p62 was recently shown to be required for aggregate formation upon proteasomal inhibition (46). To test if p62 also contributes for the non-selective apoptosis-related aggregate formation we report here, we first verified that p62 is also required for Velcade-dependent aggregate accumulation in RFP-Ova expressing 39.5 cells. Specifically, we analyzed aggregate formation in cells in which p62 expression was impaired by siRNA (Figure 5A). As seen in Figure 5B, amounts of RFP-containing aggregates, generated upon proteasomal inhibition, dramatically decreased, when p62 levels were reduced by siRNA (the efficiency of the sip62 is demonstrated in Supplementary Figure S9). Velcade-dependent aggregates co-localized with p62. Reduction of p62 levels also correlated with diminished aggregation and a more diffused RFP signal for apoptosis-mediated aggregation (Figure 5B). Notably, both Velcade-induced and apoptosis-related aggregates co-localized with p62. In further support of an involvement of p62 in apoptosis-associated aggregate formation, both p62 transcripts, as well as cellular p62 protein levels were found elevated in early apoptosis (Figure 5C). Taken together these findings underscore p62 as an important factor in apoptosis-related aggregation.

**Figure 5:**
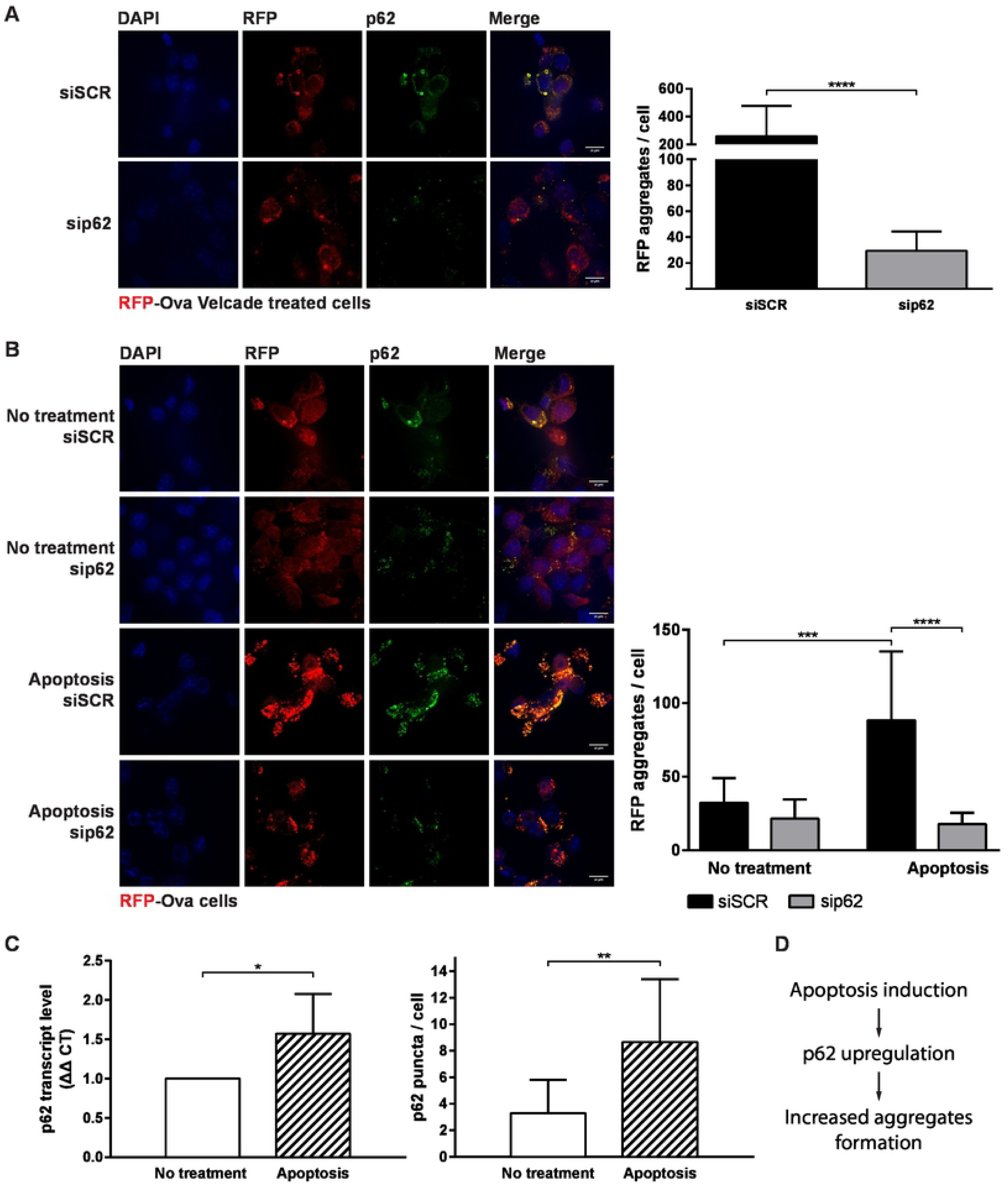
Sqstm1/p62 knockdown decreased the level of aggregate formation triggered by apoptosis. **(A)** Adherent 39.5 cells expressing RFP-Ova were transfected with either non-targeting siRNA (Scr) or with SQSTM1/p62 siRNAs separately using DharmaFect reagent. After 60 h cells were treated with 2.5µM Velcade for additional 12 hours. Next, cells were fixed, immunostained with anti-SQSTM1/p62 and DAPI, and analyzed by confocal microscopy. Scale bar: 10 μm. Quantification of RFP aggregates per cell was determined by image analysis performed by ImageJ on cells within 8 captured images (right graph). Statistical significance is p < 0.0001 and is indicated by (****). **(B)** Non treated or apoptotic 39.5 cells expressing RFP-Ova construct were transfected with either siSCR or sip62, fixed and immuno-stained with anti-SQSTM1/p62 antibody and DAPI and analyzed by confocal microscopy. Scale bar: 10 μm. Quantification of RFP aggregates per cell was determined by image analysis performed by ImageJ on cells within 12 images (right graph). Statistical significance of p < 0.001 (***) for the difference between non treated to apoptotic and p < 0.0001 (****) for the reduced phenotype in apoptotic cells. **(C)** Left graph - p62 transcript level in non-treated or apoptotic cells was measured by real time PCR. Data analysis was preformed using StepOne software v2.3. Statistical significance of p<0.05 (*), was determined using 4 independent experiments (each preformed in triplicates). Right graph – p62 protein level in non-treated or apoptotic cells analyzed by ImageJ based on fixed immunostained cells. Statistical significance of p < 0.01 (**), deduced using 12 images. **(D)** Summary scheme.

### Dendritic cells are endowed with the capacity to process polyQ aggregates and retrieve antigens for cross-presentation

To directly evaluate if aggregates can serve as source for cross-presentation, we took advantage of an Ova derivative that is fused to RFP on both termini (RFP-Ova-RFP) and was previously shown to have a very short half-life (31). This version of Ova, when delivered in apoptotic cells to mice did neither elicit T cell proliferation nor killing activity (Figure 6A and B). To test if aggregate formation could render the protein immunogenic, we equipped the RFP-Ova-RFP molecule with an aggregation-prone polyglutamine (PolyQ) encoding sequence. PolyQ stretches in proteins cause the accumulation of nonreversible deleterious aggregates and are associated with neuronal and muscular degenerative diseases (47, 48). Long PolyQ stretches are particular strong inducers of aggregation; we hence added 94 glutamine residues to the N-terminus of RFP-Ova-RFP (94Q-RFP-Ova-RFP). Immunization of mice with apoptotic cells expressing this PolyQ derivative resulted in a strong T cell proliferation and efficient CTL responses (Figure 6A and B). This suggests that promoting the aggregation of a very labile protein can serve as a strategy to induce cross-presentation and trigger cytotoxic immunity.

**Figure 6:**
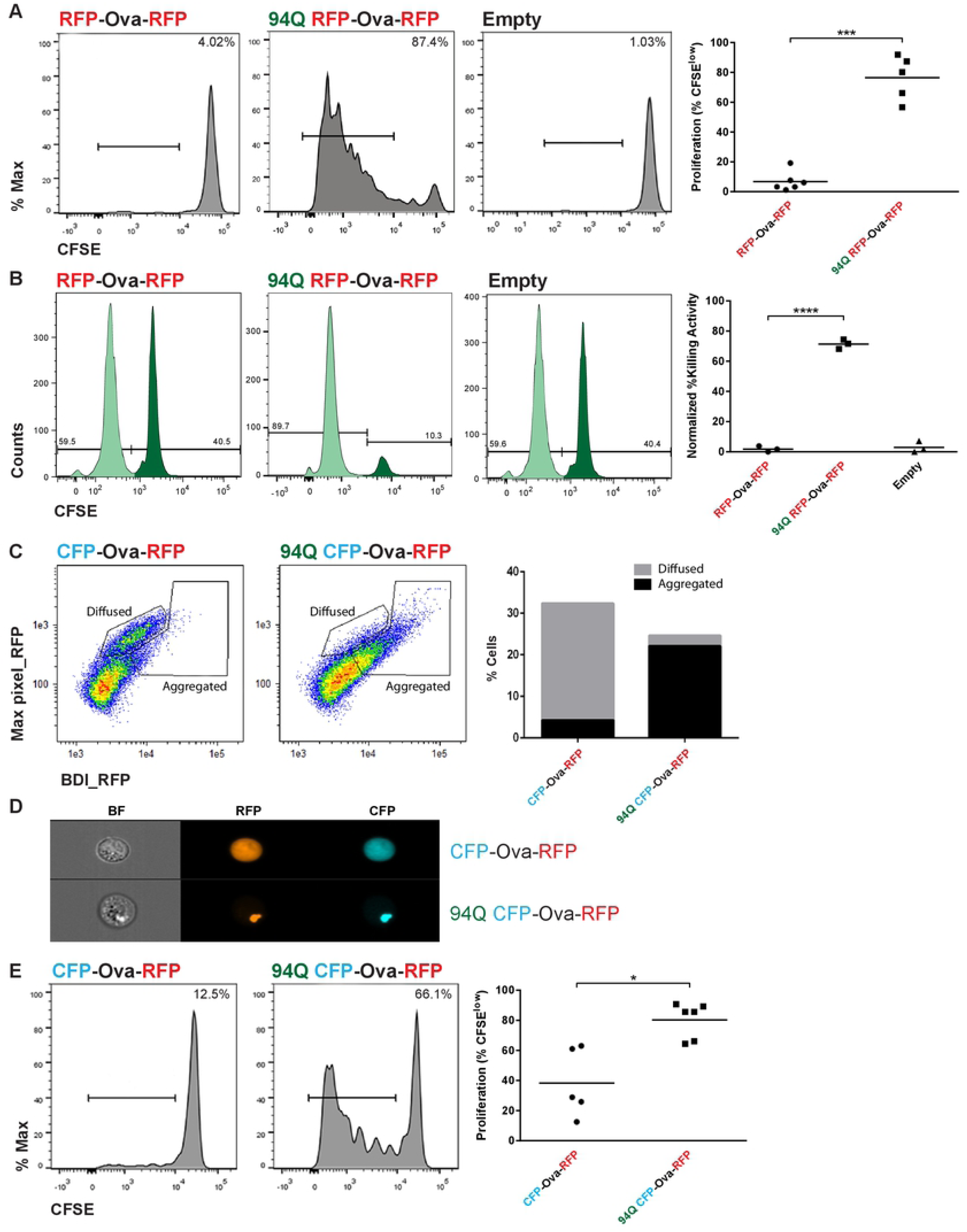
PolyQ aggregates are efficiently processed by DC and provide antigens for use in cross-presentation. **(A)** Flow cytometric analysis of lymph node and spleen cells for proliferation of CD8^+^Vα2^+^CD45.1^+^ T cell graft in mice immunized with apoptotic cells expressing RFP-Ova-RFP or 94Q RFP-Ova-RFP. Statistical significance is p < 0.001and is indicated by (***). These results are representative of two independent experiments, total n=6 (RFP-Ova-RFP) and n=5 (94Q RFP-Ova-RFP). **(B)** In vivo killing assay. Flow cytometric analysis of spleens of immunized mice injected with SIINFEKL-loaded and control target splenocytes (CD45.1^+^), labelled with high CFSE (dark green) and low CFSE (light green), respectively. Statistical significance is p < 0.0001 and is indicated by (****). These data are representative of two independent experiments, total n=6 (RFP-Ova-RFP) and n=6 (94Q RFP-Ova-RFP) **(C)** Image stream analysis of 39.5 cells expressing CFP-Ova-RFP or 94Q CFP-Ova-RFP. High BDI and high max pixel indicate aggregated RFP and CFP, while high max pixel and low BDI indicate diffused distribution of RFP and CFP. On right panel the graph represents cells percentage in each gate. **(D)** Representative image for CFP-Ova-RFP and 94Q CFP-Ova-RFP cells. **(E)** Flow cytometric analysis of lymph node and spleen cells for proliferation (CFSE dilution) of CD8^+^Vα2^+^CD45.1^+^ T cell graft in mice immunized with apoptotic cells expressing CFP-Ova-RFP or 94Q CFP-Ova-RFP, a week following removal of the tetracycline inducer from the media. Statistical significance is p < 0.01 and is indicated by (**) (representative of two independent experiments).

The fact that addition of the polyQ stretch rendered 94Q-RFP-Ova-RFP immunogenic suggests that polyQ aggregates can act as efficient protein source. Alternatively, immunogenicity could have resulted from a dynamic equilibrium between the polyQ-decorated fusion proteins *en route* to aggregation, rather than from the aggregates themselves. Moreover, some RFP-Ova-RFP degradation fragments may have re-dissolved and served as antigen source (49). Based on the fact that polyQ aggregates are endowed with extremely long-term stability, while the soluble intermediates are short-lived, we addressed this possibility by generating a TET-regulated expression system for CFP-Ova-RFP or 94Q-CFP-Ova-RFP (50). Thus, by switching off the induction of 94Q-CFP-Ova-RFP expression, soluble forms of the proteins will disappear over time, ensuring that the source for Ova epitope available for cross presentation will be predominantly, if not exclusively, aggregated protein. The different fluorescent profiles of the CFP and the RFP confirmed the integrity of the fusion proteins (Supplementary Figure S10). In addition, withdrawal of doxycycline (Dox) from the cell culture media and further growth of the cells for up to a week prior to the cross-presentation assay, ensured that no non-aggregated portions or fragments of 94Q-CFP-Ova-RFP were present in the cells. The different cellular distributions of CFP-Ova-RFP and 94Q-CFP-Ova-RFP were visualized by ImageStream. ‘Max Pixel’ analysis representing the value of highest intensity pixel within the image, and BDI analysis reflecting the intensity of localized bright spots (aggregates) confirmed that while CFP-Ova-RFP cells displayed a diffuse protein distribution (low BDI and high max pixel), 94Q-CFP-Ova-RFP appeared as typical polyQ-related aggregates (high BDI) (Figure 6C-D, Supplementary Figure S11).

Next, CFP-Ova-RFP and 94Q-CFP-Ova-RFP cells (7 days following Dox withdrawal) were triggered for apoptosis and used to challenge animals (as described in Figure 2). Significant T cell proliferation was detected only when the 94Q-CFP-Ova-RFP expressing cells were used as immunogen (Figure 6E). To further test the potency of aggregates as a source of antigens in cross presentation, we monitored the level of direct presentation in cells expressing 94Q-CFP-Ova-RFP immediately following induction. As evident, most of the cells were CFP^+^RFP^+^ and presented SIINFEKL on surface MHC-I (Supplementary Figure S12). However, 72h following Dox withdrawal, i.e. in absence of new translation, only aggregates remained in these cells, and as a consequence direct SIINFEKL presentation, was lost (Supplementary Figure S12B). In contrast, cross presentation remained efficient, as seen in the *in vivo* read out (Figure 6E). This was the case, although the cells had undergone 3-4 cell divisions and the amount of aggregates was hence 10 times diluted (Supplementary Figure 12A). This finding directly demonstrates the potency of aggregates in cross-presentation.

Taken together, these results directly demonstrate the ability of DC, and specifically cDC1, to process aggregates and support a mechanism whereby stress-related aggregation upon apoptosis is an important and potent protein source for cross-presentation.

## Discussion

Cross-presentation by DCs is critical to raise cytotoxic T cell immunity against infected and transformed cells. How the diverse pool of cellular proteins is efficiently channeled for MHC-I presentation in DCs remains incompletely understood. Here we provide evidence that apoptosis results in the formation of aggregates that can be efficiently used as antigen source for cross-presentation by DCs.

Following viral infection, pathogen-derived antigens are efficiently presented on MHC-I of infected cells (51); this is despite the fast kinetics of presentation and the fact that a significant fraction of these antigens originates from stable viral proteins. This observation led to the proposal that QC mechanisms, such as DRiPs, are a major source for direct MHC-I presentation of viral proteins and presumably other stable protein (7). To study how the epitopes contributed by DRiPs to MHCI direct presentation are captured and presented in cross-presentation, we used engineered chimeric protein constructs, which harbor a dominant ovalbumin H-2K^b^ epitope (6, 52, 53). By flipping the position of a stable globular domain juxtaposed to Ova, we created versions that are either rapidly turned-over or stable (Ova-RFP, RFP-Ova) (31). Direct MHC-I presentation of the short-lived version (Ova-RFP) was influenced by its turnover rate. Presentation of the stable protein (RFP-Ova) was a DRiP-dependent process, corroborating the notion that DRiPs are a crucial source for MHC-I presentation of epitopes originating from stable proteins. Furthermore, our findings suggest that peptides that are generated in the course of ‘translation QC’ are more efficiently processed and potentially loaded onto MHC-I. This might indicate a tighter ER retrograde transport of proteasomal products generated upon ribosomal QC degradation (54). Taken together this scenario would encode a biological bias favoring QC-associated mechanism, such as DRiPs, in MHC-I presentation of stable proteins. Notably, this would affect for instance the presentation of viral proteins, which are frequently exposed to multiple steps of translation-associated QC cycles, ensuring the proper glycosylation and assembly (55).

In the process of cross-presentation/cross-priming, presented peptides are derived from an exogenous source (25), rendering DRiPs irrelevant. In support of this notion, proteasomal degradation products of donor cells do not contribute to MHC-I cross-presentation in the recipient APCs (56). The latter is consistent with the role of cross-presentation in directing immune responses to non-self antigens, avoiding the dominating self repertoire associated with the high flux of proteasomal degradation of self short-lived proteins (SLIPs) (57) and QC rejects.

In view of the importance of repertoire overlap of direct and cross-presentation for effective immune responses (30), it is unclear how extremely short-lived entities, such as DRiPs, can be captured by DCs, as only limited amounts of them are available for cell-to-cell transfer. Our data suggest that global nonselective aggregation is a significant biological process that enables capturing the steady state abundance of the cellular stable proteome engulfed by DCs, in a manner conducive for presentation.

We propose that apoptosis entails global protein aggregation, which ensures that the cross-presented protein spectrum mirrors the contribution of DRiPs and QC-related epitopes in MHC-I direct presentation. Our findings suggest that this mechanism enforces representation of antigens originating from stable proteins at the expense of very short-lived proteins, thereby directing immune responses towards relevant antigens, including viral proteins in the case of infected cells. In addition, involvement of protein aggregation is consistent with earlier studies demonstrating that stable proteins are the major contributors to cross-presentation (56, 58–61). This is also in agreement with the proposed role of heat shock proteins (HSPs) in cross-presentation (62), as these serve as potential promoters and stabilizers of aggregates. In fact, molecular chaperones were implicated as a mechanism to facilitate the transfer of antigenic peptides derived from stable proteins to cross-presentation (63, 64). In support of this model, various HSPs, including hsp70, hsp90, and grp94 gp96, isolated from tumors, as well as infected or transfected cells yield antigenic peptides (63). Notably, the function of aggregates in cross-presentation also highlights a potential role for autophagy in the processing of aggregates. This notion is further supported by the role of p62 in apoptosis related aggregation, as p62 may serve for targeting of the aggregates to autophagy upon intake by DCs.

To directly assess if aggregates can serve as a significant source for cross-presentation, we utilized polyQ to drive aggregation of our proteins of interest. PolyQ expansion-related pathologies result from faulty proteins that harbor abnormally long stretches of glutamine residues. The extent of aggregation and the severity of the related disease directly correlate with the length of the polyQ stretch, where 94Q is in the very upper range of the scale (65, 66). Using this system we demonstrated directly that aggregates serve as efficient source for antigens in cross-presentation and that DCs are endowed with a robust ability to process even extreme aggregates.

Notably, global aggregation was not observed in heat-induced necrotic cell death suggesting that aggregates are not an exclusive source of antigens for cross-presentation, but that there are multiple ways for APCs to acquire intact antigens from their environment. As phagocytes, APCs efficiently internalize dying cells and their debris, as well as shed exosomes, all of which contain intact antigens (28, 67–69). It has also been reported that DCs can actively ingest pieces of antigen-bearing cells (70). In support of this notion, intact protein antigens, both in particulate form and, less efficiently, in soluble form, can be cross-presented on MHC class I by professional APC, at least in *in vitro* systems (60). In this respect, intake of aggregates as cellular debris or within exosomes (71) would be advantageous as the mass / volume ratio of relevant material to be used in cross-presentation is higher compared with intake of soluble proteins in more diluted form. Either way, the fact that cross-presentation relies on both lysosomal and proteasomal processing (25) is beneficial, as lysosomes were shown to process polyQ-related aggregates. This leaves the trafficking of the aggregates to the lysosome as a potentially limiting factor (72), and highlights the relevance of p62 as a potential lysosomal targeting factor of apoptosis-related aggregates in DC.

Interestingly, a functional role for aggregates in DC, in the context of APC capacity, was already noted in prior studies (73–75). In these studies, DC aggresome-like induced structures (DALIS) were found to facilitate MHC-I cross-presentation of exogenous (non-self) antigens by diverting self-ubiquitinated proteins (including self-DRiPs) to transient aggregates. Although related, these structures are distinct from the aggregates in the current study, which are rather stable, produced in the donor cells and in contrast to DALIS serve as major antigen source for cross-presentation. Taken together with our current findings, we conclude that DC have evolved to use aggregation both to reduce the self MHC-I antigens load upon immunological challenge, as well as to select relevant antigens in target cells for cross-presentation.

Future detailed studies of the mechanisms that control aggregation in the donor cells and de-aggregation and processing in the APC are important to understand the pathways underlying the generation of CTL-mediated immune responses and exploit them for therapeutic vaccination strategies. Furthermore, the robust and unique ability of DC to process polyQ aggregates, even when aggregation is driven by a long polyQ stretch (94Q), as illustrated here *in vivo,* demonstrated for the first time an efficient physiological capacity of DC to process polyQ aggregates, probably relying on lysosomal catabolism (76). This might suggest a potential role for the immune system in clearing aggregates that should be further studied and potentially could be exploited for therapy.

## Methods

### Mice

C57BL/6 (CD45.2) WT mice were purchased from Harlan (Rehovot, Israel). OT-I T cell receptor (TCR) transgenic mice (33) were bred in the animal facility of the Weizmann Institute. BM chimeras were generated using WT and BatF3 KO BM (kindly provided by Kenneth Murphy) (18). 8 weeks old female recipient animals (CD45.1/1) were lethally irradiated (950 rad) and reconstituted with female donor BM by i.v. injection. Mice were kept under Ciproxin (Bayer) antibiotics for 10 consecutive days and BM chimeras were analysed 8 weeks after transfer. All mice were bred and maintained in specific pathogen-free (SPF) animal facilities at the Weizmann Institute of Science. Experiments were approved by an Institutional Animal Care Committee (IACUC) in accordance with international guidelines.

### Cell lines

39.5, an H-2K^b^ D122 transfected cell line derived from 3LL Lewis lung carcinoma (kindly provided by Lea Eisenbach) (77), were cultured in DMEM supplemented with 10% fetal bovine serum, 2mM glutamine, 50 units/ml Penicillin, 50µg/ml streptomycin, 1x nonessential amino acids, 1 mM sodium pyruvate and 0.5 mg/ml Geneticin. EL4 (mouse thymoma cells) were cultured in RPMI 1640 medium supplemented with 10% fetal bovine serum, 2mM glutamine, 50 units/ml Penicillin, 50µg/ml streptomycin, 1x nonessential amino acids and 1 mM sodium pyruvate. All cells were grown at 37°C and 5% CO_2_.

### Generation of EL4 and 39.5 cells expressing ovalbumin variants fused to RFP

Ova fused to RFP was cloned into pMIG, a retroviral expression vector, which harbors an IRES-GFP element. Viruses were generated in 293T cells by triple transfection with Env and Gag-pol encoding vectors, as described (78). Transduced EL4 or 39.5 cells were sorted to GFP-positive to 100% purity by FACS (Aria, BD) and cultured similarly to parental EL4 cells. Poly glutamine (94Q) fused Ova derivatives were cloned into pTRE-tight, a mammalian expression vector. 39.5 cells were co-transfected with pCMV and pTRE-CFP-Ova-RFP or 94QCFP-Ova-RFP in the ratio of 1:4 using the JetPEI reagent (company). CFP and RFP positive cells were sorted by FACS (Aria, BD) and cultured similarly to the parental 39.5 line.

### Monitoring the levels of H-2K^b^/SIINFEKL complex on the EL4 cell surface

10^6^ EL4 cells of interest were washed with FACS washing buffer (PBS, 0.5% BSA, 0.1%NaN_3_) and incubated for 60 min on ice with 25D1.16, a mouse monoclonal antibody, which recognizes the H-2K^b^/SIINFEKL complex (32). Cells were then washed three times with the buffer and incubated on ice with Alexa Flour 647-conjugated anti-mouse polyclonal antibody for 60 min (Molecular Probes). Consequently, the cells were resuspended in PBS and analyzed by FACS (BD biosciences). Mean fluorescence intensity (MFI) was calculated, using FACSdiva software (BD biosciences). EL4 cells were treated with 5µM Velcade (Bortezomib, Selleck Chemicals, 179324-69-7) to inhibit the proteasome activity, and 50µg/ml chyclohexamide (CHX) (Sigma, C7698-1G) to inhibit the ribosome activity.

### Peptide stripping by acid wash

To remove MHC class I bound peptides from the cell surface, cells were washed with citrate phosphate buffer, pH 3.0 (0.263 M citric acid and 0.123 M disodium phosphate), as described (79). Briefly, cells were washed twice with PBS and then incubated for 1 minute with 0.3 ml citrate phosphate buffer at room temperature. Afterwards, cells were washed twice with RPMI 1640 medium, supplemented with 10% fetal bovine serum, 2 mM Glutamine, 50 units/ml Penicillin, 50 µg/ml Streptomycin, 1x nonessential amino acids and 1 mM Sodium Pyruvate. Subsequently, cells were resuspended in fresh medium and analyzed by FACS, or further incubated with the relevant reagents.

### B3Z T cell activation assay

Sixty thousand B3Z cells, a *LacZ*-inducible CD8^+^ T cell hybridoma specific for the SIINFEKL/H-2K^b^ complex (80), were incubated with serial dilutions of the respective 39.5 cells in flat-bottom 96-well plates for 12 h. LacZ activity was detected by incubating with 9 mM MgCl2, 0.125% NP-40, 0.3 mM chlorophenol red β-D galactopyranoside (CPRG) in PBS for 30 minutes at 37°C. OD595 was determined using a Varioskan Microplate reader.

### Adoptive cell transfer

CD8^+^ T cells (isolated from 10 weeks old female OT-I mice) were positively selected from CD45.1^+^ donor spleens using anti-CD8α beads (Miltenyi biotech). OT-I T cell proliferation was assessed by staining with 5µM carboxyfluorescein diacetate succimidyl ester (CFSE; (34)) for 8 min at RT in RPMI medium without serum. CFSE labeling was quenched by adding two volumes of cold FCS and 5 min incubation on ice. Cells were washed in RPMI medium and 1X10^6^ cells were transferred intravenously to recipient naïve C57BL/6 WT mice. Probe dilution was quantified 5 days later using flow cytometry.

### Apoptosis induction and antigen administration

39.5 cells expressing either ovalbumin variants fused to RFP or no vector (Empty cells) were incubated for 12 hours with 500 µM Hydroxyurea (SIGMA, H8627-1G) to induce cell synchronization. Apoptosis was induced with 100 µM Etoposide (SIGMA, E1383-25MG) for an additional 12 hours. Thereafter, the cells were washed with PBS, and 1X10^6^ apoptotic cells were injected intravenously to the C57BL/6 WT recipient mice, 24 hours following OT-I CD8^+^ T cells transfer.

### Necrosis induction and AnnexinV/DAPI labeling

39.5 cells expressing ovalbumin variants fused to RFP were collected and incubated for 5 minutes at 65°C. Then cells were washed with PBS and incubated for 15 min at room tempature with 510 bright field annexinV; 5 µl in 100 µl binding buffer (50 mM Hepes, 700 mM NaCl, 12.5 mM CaCl2, pH 7.4). Following an additional wash with PBS Draq5 and DAPI were added to the cells which were analyzed by ImageStream.

### In vivo cytotoxicity assay

The *in vivo* killing assay was performed as described before (35). 7 days following apoptotic cells injection, splenocytes were isolated from WT CD45.1^+^ mice. Half of the cells were pulsed with 10µg/ml SIINFEKL peptide for 2h at 37°C. Unpulsed and pulsed cells were stained with 0.5 or 5 µM CFSE, respectively, and a 1:1 mixture of the cells was transferred to immunized C57BL/6 WT mice. The differential clearance of both target cell populations was evaluated using flow cytometry 24 hours later. Specific killing was defined as 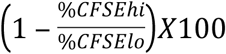. This value was normalized to the average specific killing of control mice, immunized with apoptotic cells harboring empty vector.

### Western Blot analysis

Total cellular proteins were prepared in NP-40 buffer (25 mM Tris, pH 7.4, 150 mM NaCl, 1% IGEPAL CA-630 [Sigma, I8896]), the extracts were centrifuged for 20 min at 21,000 x g and lysates were separated to supernatant and pellet. Sup protein concentration was determined using Pierce BCA protein Assay Kit (Thermo scientific, 23225). Twenty µg protein was loaded from sup and the corresponding amount was loaded from pellet. Proteins were separated on gradient 7%-15% SDS-PAGE gel. The membrane was blocked in 10% BSA for 30min at room temperature. After blocking, the membrane was incubated with appropriate primary antibody at 4°C overnight or at room temperature for 3 h. Then the membrane was washed with PBS-TWEEN20 (PBS supplemented with 0.5% TWEEN-20, [Sigma, P1379]), and incubated with the secondary antibody (goat-anti-mouse or goat-anti-rabbit) at room temperature for 45 min. Finally, the membrane was washed 3 times and the specific proteins were visualized using the Enhanced ChemiLuminescence (ECL) detection system (Biological Industries 20-500-120). Antibodies used for western blot in this study: home-made rat α-RFP antibody, mouse α-actin (Santa cruze sc-376421) and rabbit α-lamin B1 (Cell Signaling #12586).

### Multispectral imaging flow-cytometry (Imagestream) analysis

Untreated cells and cells following induction of apoptosis were imaged using multispectral imaging flow cytometry (ImageStreamX flow cytometer; Amnis Corp). For multispectral imaging flow cytometry, approximately 3×10^4^ cells were collected from each sample and data were analyzed using image analysis software (IDEAS 6.0; Amnis Corp). Images were compensated for fluorescent dye overlap by using single-stain controls. Cells were gated for single cells using the area and aspect ratio features, and for focused cells using the Gradient RMS feature, as previously described (43). Bright detail intensity (BDI) computes the intensity of localized bright spots within the masked area in the image. The local background around the spots is removed before the intensity computation. Area Threshold 50% calculates the area of the 50% highest intensity pixels within the image. Max Pixel is the value of highest intensity pixel within an image, ranging from 0 to 4095.

### siRNA transfection

For siRNA, adherent 39.5 cells was transfected using DharmaFect1 (Dharmacon, T-2001-04) according to the manufacturer’s instructions with different siRNA SMARTpools (50 nM each) targeting *SQSTM1/p62* (Thermo scientific L-047628-01-0005) and scrambled nontargeting siRNAs control (Thermo scientific D-001810-10-05). Experiments were performed 72 h after transfection. Total cell extracts were prepared using NP-40 extraction buffer (25 mM Tris, pH 7.4, 150 mM NaCl, 1% IGEPAL CA-630 [Sigma, I8896]) with a protease inhibitors mixture. The knockdown effect was determined by western blot analysis.

### Tissue-culture Immunostaining

Adherent cells (39.5) were grown on lysine coated 24 well plate. The cells were fixed with 100% Methanol for 7 min and blocked with PBS containing 0.1% Triton and 20% normal horse serum for 90 min at room temperature. For immunostaining, the cells were incubated for 1h at room temperature with anti-SQSTM1/p62 monoclonal antibody (H00008878, abnova) diluted in the blocking solution. Cells were then washed three times with PBS and stained for 30 min at room temperature with suitable secondary antibody. This was followed by 5 min of DAPI (4,6-diamidino-2-phenylindole dihydrochloride). The cells were viewed under Nikon fluorescence imaging microscope.

## Acknowledgments

We thank Lea Eisenbach and Adi Sharbi Yunger (Weizmann Institute of Science) for providing advice and critical reagents. We thank K. Murphy for provision of BATF3 BM. The Jung lab was supported by the European Research Council (AdvERC 340345) and the Israeli Science Foundation (887/11). A.N. research was supported by the Israeli Science Foundation (2038/17).

## Autor Contribution

S.T.C. performed and analyzed experiments. D.B. provided advices and performed experiments. S.P, Y.W. and C.C., provided advice and technical assistance. Z.P. helped perform the Image stream assay and analysis. R.K. provided technical assistance. B.T helped writing the manuscript. S.T.C., S.J. and A.N. planned the project, designed the experiments and wrote the manuscript.

